# Exploiting Four-way Phase-encoding Benefits for Robust Detection and Correction of EPI Artifacts: Application to Residual Ghosts in Diffusion MRI

**DOI:** 10.1101/2024.12.11.627992

**Authors:** Anh Thai, Lin Ching Chang, Carlo Pierpaoli, M. Okan Irfanoglu

## Abstract

**Purpose:** To propose and develop an image processing-based methodology for detecting and correcting echo planar imaging (EPI) artifact that employs data from 4-way phase-encoding acquisitions.

**Approach:** Previous studies have demonstrated that acquiring images with four different phase-encoding directions can improve the reproducibility of diffusion derived quantitative maps while retaining the same acquisition time. This improvement is achieved by averaging across phase-encoding directions (PEDs) to reduce the impact of EPI artifacts. Building on this principle, the proposed method further improves this 4-way encoding approach by leveraging the properties of signal distributions to detect artifactual regions. Additionally, the method can be tailored for specific artifact manifestations by considering their localization due to underlying acquisition parameters. The proposed method is applied to and validated using both simulated data and in-vivo diffusion MRI data affected by residual ghost artifacts.

**Results:** Simulations with known ground-truth images demonstrated high artifact detection accuracy, achieving a Dice score of 0.91 for reconstructions without parallel imaging. In the in-vivo dataset, the method also improved longitudinal reproducibility, reducing variability by 30% in ghost-affected regions.

**Conclusion:** The proposed correction method effectively detected and corrected residual ghost artifacts without the need of any additional k-space data. This retrospective approach can be directly integrated into existing processing pipelines to further improve the quality of EPI images and enhancing image quality in studies that utilize 4-way PEDs acquisition.

## 1. Introduction

Echo planar imaging (EPI) distortions and artifacts have significant impacts on diffusion MRI (dMRI)-derived scalar maps (1–3), tractography (4,5), and the reproducibility of multi-center study findings (6). Correcting EPI distortions generally requires additional data, such as field maps (7,8) or anatomical images for elastic registration (9,10). In recent years, the use of multiple phase-encoding directions (PEDs) during acquisition (11) has emerged as a promising approach to improve image quality, as it shows superior performance in EPI distortion correction compared to the alternative methods. Despite evidence supporting the advantages of acquiring all dMRI volumes with more than one PED (12), the most common approach is to acquire the reversed phase-encoding data only on the non-diffusion weighted, *b* = 0 *s*/*mm*^2^ volumes (7,13).

In a previous study, Irfanoglu et al. (12) evaluated the reproducibility of different acquisition schemes, comparing single versus multiple phase-encoding directions (PEDs), while ensuring that each scheme had approximately the same acquisition time. Among these, the scheme using four different PEDs such as anterior-posterior (AP), posterior-anterior (PA), left-right (LR), and right-left (RL), showed improvements in reproducibility, particularly by enhancing distortion correction and mitigating the impact of ghost artifacts. This artifact remediation was achieved because, in EPI, ghost artifacts manifest at different spatial positions depending on the PED. For instance, ghosts of the eyes might overlap with the brain when the images are acquired with AP or PA encoding, whereas when acquired with LR or RL directions, eye ghosts are laterally positioned outside of the brain regions. This ensures that while artifacts may still occur, the use of four phase-encoding directions guarantees that for a given region of interest, at least two of the PEDs should not be affected by the same artifact. Thus, with four-way phase-encoding, simple methods such as data concatenation or averaging (as performed by Irfanoglu et al. (12)) can effectively reduce artifacts by up to two-fold.

Inspired by this previous study, rather than relying solely on this artifact mitigation strategy, we propose a pipeline that delves deeper into existing data to further improve the reproducibility by detecting the artifacts across phase-encoding images based on their signal characteristics, eliminating/reducing their impact, and retrieving the correct information from other phase-encoding directions. This approach leverages the “good data” from unaffected PEDs and outperform the averaging method from the previous study. The proposed EPI artifact correction strategy is adaptable to various artifact types that vary in spatial location based on PED, including ghost artifacts, chemical shifts, parallel imaging artifacts, etc. In this study, we demonstrate the proposed framework on residual ghost artifacts in dMRI to evaluate its effectiveness. We define “residual artifacts” as artifacts that remain after conventional EPI phase-correction (ghost artifact removal) has been applied by the scanner.

Several approaches exist to remove Nyquist ghost artifacts in EPI data. The most common strategy is to estimate the linear phase differences between even and odd echoes by using a calibration scan (8,14–16). More advanced techniques, such as Navigator-based phase correction strategies (17–20), reference-free methods (21–28), and deep learning-based methods are also frequently used (29–31). These methods are ideally applied at the acquisition stage, as they often rely on access to raw k-space data or calibration images. However, once initial corrections are applied and images have been fully reconstructed, residual ghost artifacts are often observed (18,32,33), making further correction challenging, as end-user don’t have access to raw data. In addition, in-plane parallel imaging techniques can exacerbate the effect of such artifacts (34–36).

The manuscript is organized as follows: Sec. 2.1 describes how information from four-way PED data can be utilized to detect signal artifacts. Sec. 2.1.2 provides a specific strategy, focusing on residual ghost artifacts. Sec. 2.3 and 2.4 detail our validation strategies using both simulated and in-vivo data, followed by the presentation of our results. Finally, we conclude the manuscript with discussion of the strengths and disadvantages of this approach, while describing how other modalities and artifact types can benefit from similar strategies.

## 2. Methods

**Figure 1** showcases an example 4-way dataset suffering with artifact localized at different positions across native-space PED images (two top rows). To capitalize on the spatial variability of residual ghost artifacts across PEDs for artifacts detection and correction, the methodological framework proposed here assumes that all necessary image corrections have been performed during acquisition, reconstruction (on-site), and DWI processing (off-site, see Sec. 2.4.1). For example, **Fig. 1** (two bottom rows) also showcases the same data after processing, where the effects of EPI distortions, motion artifacts, noise and physiological variation are minimized, leaving the residual artifacts as the predominant differences. These residual artifacts manifest at different spatial positions across images acquired with various PEDs, as highlighted by yellow arrows.

**Fig. 1.**
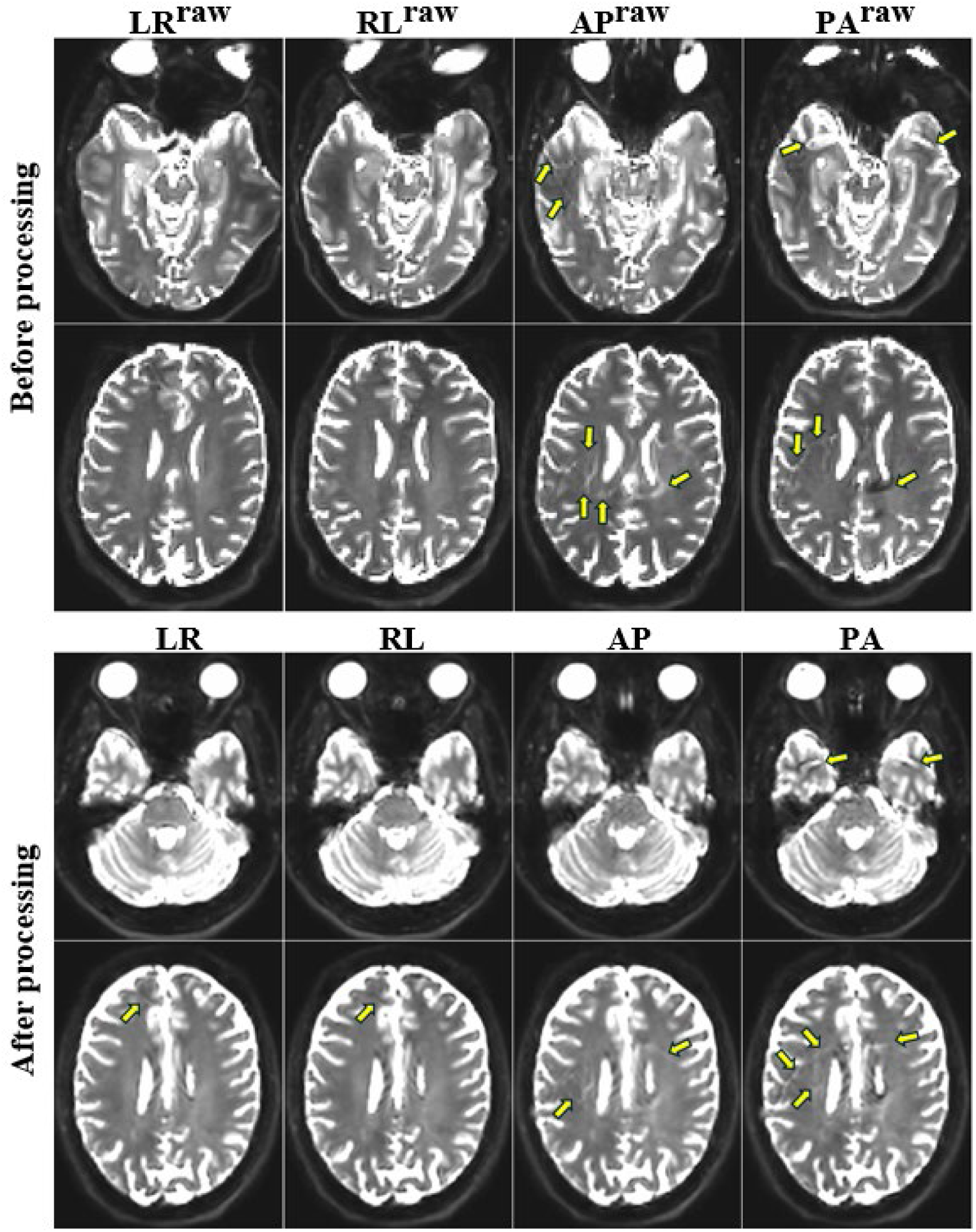
Manifestation of residual ghost artifacts before and after processing of non-diffusion weighted *b* = 0 *s*/*mm*^2^images across four different PEDs. Two slices from the *b* = 0 *s*/*mm*^2^images of a representative subject acquired with four different PEDs: LR^raw^ (left-right), RL^raw^ (right-left), AP^raw^ (anterior-posterior), and PA^raw^ (posterior-anterior) are selected. In these raw, unprocessed images, which suffer from EPI distortions, the yellow arrows highlight the residual artifacts with varying intensities, particularly in the AP and PA images. After processing to correct typical dMRI distortions, the images are aligned to the structural image, resulting in four nearly identical images, denoted as LR, RL, AP and PA. Despite the corrections, subtle residual artifacts persist as indicated by the yellow arrows.

To guide the detection and correction process, this framework operates as follows:

- EPI artifacts, including residual ghosts, appear at different spatial positions among PEDs and can be present in single, multiple or all PED images as either hyper- or hypo-intensities. These signal outliers result in larger signal variances among PEDs which can be detected through a statistical method.
- Residual ghost synthetic map: with real in-vivo data, signal variations can also arise from other imperfections such as inaccurate processing, physiological noise, etc., resulting in false-positive detections. To improve the detection and isolate residual ghost artifacts, a model is proposed to mimic the actual residual artifacts in the images. Using this model, a synthetic ghost map is generated and segmented with Otsu method (37), to automatically isolate regions likely affected by artifacts.

The results are combined to produce comprehensive residual ghost artifact detection across the four PEDs by mitigating the detected artifacts using two different correction methods.

### 2.1 Artifact detection methods

#### 2.1.1 Variance-based analysis (VBA)

Our analysis focuses on estimating how artifacts across different PEDs influence the overall variance observed in the images. Let *Y* be a set of all samples from PEDs, defined as *Y* = {*V*_*LR*_, *V*_*RL*_, *V*_*AP*_, *V*_*PA*_}, where each *V*_*i*_ is the set of voxel intensities drawn from a [3 × 3] neighborhood window from all *b* = 0 volumes of each corresponding PED. To analyze the local variance, we apply the law of total variance, such as:

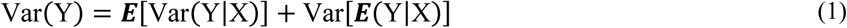

This formula breaks down the total variance into components that represent the average variance within each PED and the variation between each PED. Equation (1) can be adapted as follows for voxel-wise analysis:

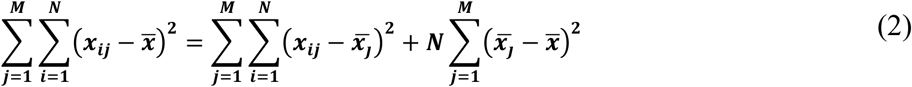

Here, the mean voxel intensity for each PED is denoted as 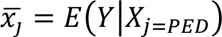, and individual voxel intensities within each PED are represented by *x*_*ij*_. N and M are the number of data points of each *V*_*i*_ and the number of PEDs, respectively. N remains constant across PEDs since each PED has the same number of *b* = 0 volumes. Therefore, the variance of each PED group relative to the overall mean, 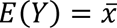, is quantified and denoted as ***X***_*j*_, which can be expressed as:

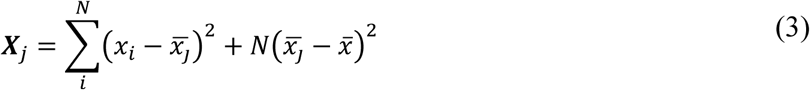

Thus, the total variance can be computed from the summation:

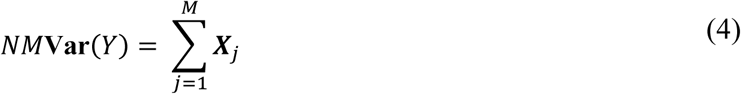

Therefore, the relative variance contribution of each PED to the total variance can be expressed as a percentage:

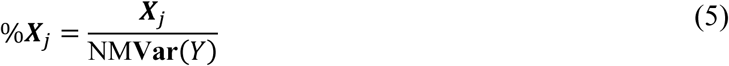

In the absence of significant artifacts, the variance contribution of each of the four distinct PEDs would be approximately 25%. If the variance contribution (i.e., the VBA score) of a PED significantly exceeds 25%, this indicates an imbalance in voxel intensities across PEDs. Therefore, that voxel can be flagged as containing an artifact. In this work, a heuristic threshold of 30% was selected for detection.

#### 2.1.2 Artifact mask generation by residual ghost synthetic image

To further improve detection data and reduce false positive, one can focus solely on voxels that can be impacted by the artifact in consideration based on acquisition parameters as we are considering the residual ghost artifacts as illustration case, in this section we will demonstrate how this can be achieved for such artifacts.

Because ghost artifacts originated from the errors between odd and even echoes, the separation of odd and even in any acquisitions will impact the strength and shift of artifacts. As the phase parameters are not estimated accurately, discrepancies arise, denoted as 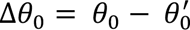 and 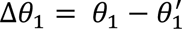, where θ^′^ are the estimated parameters and θ_0_ and θ_1_ are the true parameters of the main field offset and constant echo delay respectively, as described by Buonocore et al. (38). Ghost elimination with linear phase methods operated in the hybrid domain *H*(*x*, *k*_*y*_), where a one-dimensional inverse Fourier Transform, denoted as 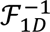 was applied along the readout direction of k-space. In this domain, the full correction is represented as:

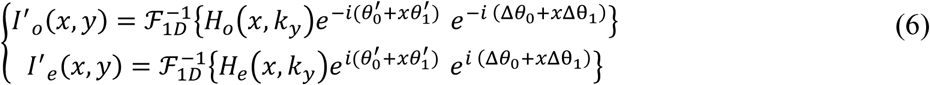

Here, *H*_*o*_ and *H*_*e*_ represent hybrid domains of ideal odd and even lines in k-space respectively. Another one-dimensional Fourier transform is applied along the PED, resulting in 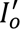 and 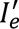 which are images of odd and even lines after phase correction, which contain residual artifacts. Assuming that the last exponent terms originated from an incomplete correction, we hypothesized that Δθ should be sufficiently small. Then, by Taylor’s series expansion, Eqn. (6) can be expressed as:

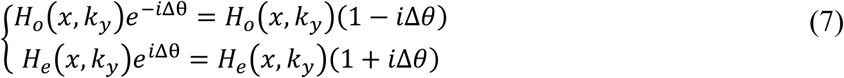

Therefore, the final image with the residual artifacts becomes:

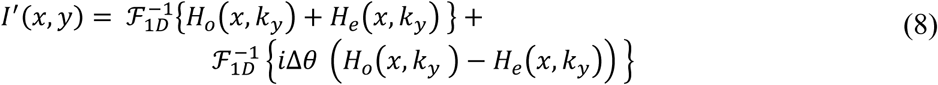

The term involving Δθ(*x*), which delineates the difference between the actual phase errors and their residuals, functions as a linear high-pass-filter (HPF) (22). This HPF correlates with the readout direction, which attenuates lower frequencies, and intensifies linearly as frequency increases. Therefore, due to the linearity property of Fourier transform, the synthetic artifact image can be simply expressed as:

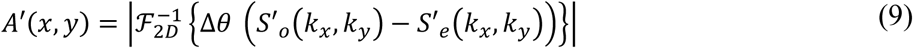

The HPF is of the form Δθ = |(*x*. Δ*k*_*x*_)|, treating positive and negative frequencies equally, attenuating lower frequencies without introducing any asymmetry or bias (39,40). Here, *S*′_*o*_(*k*_*x*_, *k*_*y*_) and *S*′_*e*_(*k*_*x*_, *k*_*y*_) are odd and even lines in virtual k-space which is a two-dimensional Fourier transform (ℱ_2*D*_) of reconstructed image *I*′(*x*, *y*). This difference between odd and even lines reflects the ghost artifact’s shifting properties (15). The synthetic ghost image *A*′(*x*, *y*) is then segmented using an automatic Otsu thresholding method (37) and morphological closing operation to produce a binary mask of areas of interest, 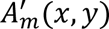, as described in **Fig. 2**.

**Fig. 2.**
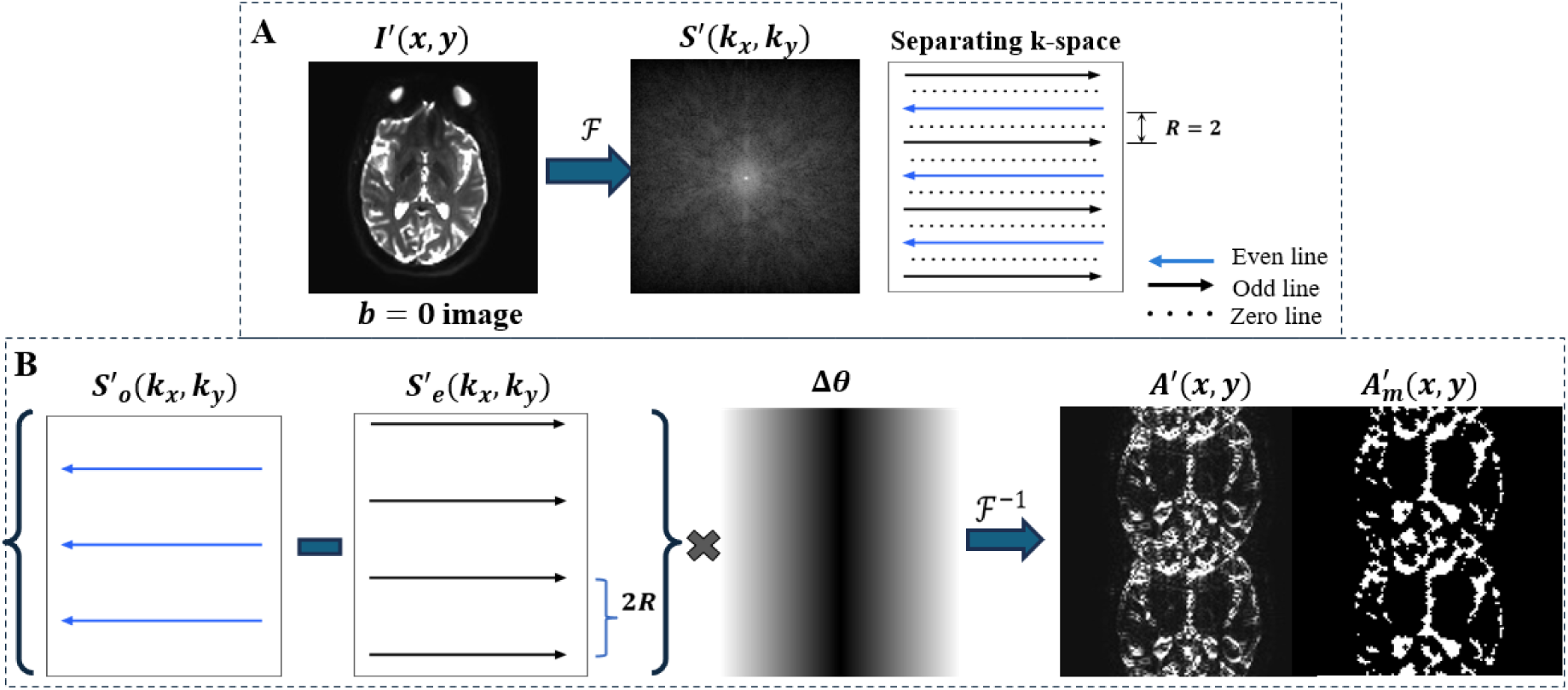
Generation of a synthetic ghost image with R=2 and PED in AP/PA directions. In Panel A, the image ***I*^′^(*x*, *y*)** is 2D Fourier transformed into virtual k-space, ***S*^′^(*k*_*x*_, *k*_*y*_)**. Odd, even, and zero lines are separated based on the undersampling factor R along the PED (AP or PA). In Panel B, the difference between odd and even lines is computed, and the HPF ***Δθ*** is applied, followed by an inverse 2D Fourier transform to produce a synthetic residual ghost image, ***A*′(*x*, *y*)**. Otsu thresholding and morphological closing operations are subsequently employed to generate the synthetic ghost map, 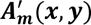.

### 2.2 Correction methods

We deploy two different correction approaches, CorrV1 and CorrV2:

#### 2.2.1 Artifact voxel removal and averaging (CorrV1)

Our first approach aims to generate an artifact-free image by combining data from the four PEDs. We begin by identifying and removing artifactual voxels, specifically those exceeding a heuristic threshold (i.e., 30%). Following removal, we average the corresponding voxels from the other unaffected PEDs. The resulting corrected image is denoted as *C*_1_. This method proves particularly effective in eliminating the influence of prominent artifacts, thus yielding a cleaner composite image.

#### 2.2.2 Averaging using synthetic ghost images and VBA scores as weights (CorrV2)

Our second approach employs a weighted average scheme. The weights are determined based on the presence and degree of artifacts, as captured by synthetic ghosts and computed VBA scores. These weights are calculated using the formula 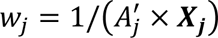, where 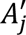 represents the artifact influence of the *j^th^* PED image as measured from the synthetic ghost image and ***X***_*j*_ represents the variance contribution of that PED image in percentage. The corrected image is then obtained by applying the following equation:

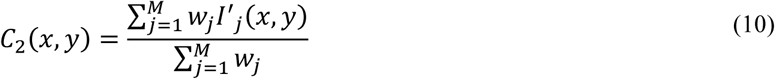

In this equation, 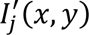 is the original image intensity at location (*x*, *y*) of the φ^th^ PED image. This method ensures that original images with higher artifact influence, as revealed by synthetic ghost images and VBA, are assigned less weight in the computation of the corrected image intensity. Therefore, this reduces their impact on the final image, denoted as *C*_2_. In cases where no artifacts are present across PEDs, (i.e. *w*_*j*_ → ∞, ∀*j*), we set *w*_*j*_= 1 for all *j*, thus the equation above become a standard average of the original images.

### 2.3 Evaluation using simulated data

To validate the proposed methods, we generated a series of simulated data. We utilized a T2-weighted (T2W) 2D fast spin-echo image from an existing in vivo dataset, which served as an artifact-free ideal image, denoted as *I*(*x*, *y*). Residual ghost artifacts, *A*(*x*, *y*), were generated using Equation (9), with the separation of odd (*S*_*o*_) and even (*S*_*e*_) lines from 2D Fourier transform of *I*(*x*, *y*) along the PED. Additionally, a mask *A*_*m*_(*x*, *y*) was generated based on the proposed thresholding to isolate the artifact structure, serving as a ground truth label for artifact detection. The artifact image introduced to the ideal image was ℳ(*A*(*x*, *y*)) = *A*_*m*_(*x*, *y*)*A*(*x*, *y*). This allowed us to focus on the voxels affected by ghost artifacts rather than additional signal, and noise variations that can be introduced by high pass filtered images *A*(*x*, *y*).

A control variable, Ω, was introduced and randomized within the range [-1.5, 1.5] to govern the severity of residual artifacts in the generated images, allowing us to simulate a broad spectrum of arbitrary artifact strengths typically encountered in practice. Artifacts generated with an Ω value within [−0.5, 0.5] were considered negligible. The generated aliased images can be delineated as shown in Eqn.(11) below, and in **Fig. S1**:

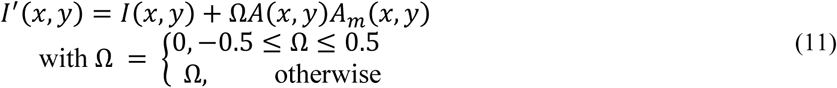

Two distinct datasets were simulated to assess the method’s performance under different imaging conditions. The first one was designed to mimic EPI images with a fully sampled k-space, labeled as R=1. In this setup, mismatches between the even and odd k-space lines caused residual ghost artifacts to appear at the N/2 position along the PED of the image. The second design was tailored to simulate a parallel imaging scenario with an acceleration factor of 2, denoted as R=2, resulting in residual ghost artifacts appearing at N/4 position.

To simulate all possible scenarios that can occur when residual artifacts exist across PEDs, four cases were generated for both R=1 and R=2 datasets: in Case 1, only one arbitrary image suffered from residual ghosts mimicking isolated artifacts when scanner corrections performed well for three of the four PEDs. For Cases 2, 3 and 4, two, three, and four images across all PEDs were synthesized with artifacts, respectively. For each case, 100 samples were generated with differing artifactual PED combinations and varying degrees of artifact strength Ω. Ghost artifacts contributed positively or negatively to the signal depending on the polarity of the phase encoding direction (14,38).

**Detection assessment:** the performance of our artifact detection method was assessed by comparing its results with the groundtruth images *A*_*m*_(refer to **Fig. S1** and **Fig. S2**). Various performance metrics including Dice similarity coefficients, Recall (true positive rate), and Precision were employed to measure the accuracy of detection method. These metrics were calculated based on counts of true-positive (TP), true-negative (TN) false-positive (FP), and false negative (FN), as shown in the following formulars:

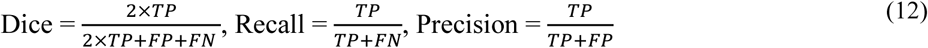

**Correction assessment:** the efficacy of our correction methods was evaluated using two metrics: the Structural Similarity Index (SSIM)(41) and Absolute-Percentage Errors (%*Err*) as shown in Equation (12). SSIM is used to measure the visual similarity between the images, which is essential for assessing the perceptual quality of the correction. The %*Err* quantifies the percentage differences between the corrected images, 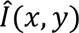 with the ideal image, providing a numerical measure of correction quality.

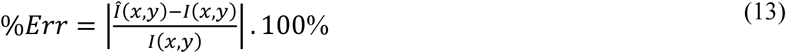

These metrics were applied to compare both the mean image, derived from averaging all PED images, denoted as *C*_0_ and our corrected images, each with the ideal image. This comparison establishes a baseline to evaluate whether our correction methods improve upon the standard averaging technique.

### 2.4 Evaluation using vivo data

The dMRI data used in this study was obtained from prior research by Irfanoglu et al. (12), which includes five longitudinal scans from seven healthy subjects acquired over six months using a Philips Achieva 3T MRI system. The study was carried out under the Institutional Review Board approved protocols and all volunteers provided informed consent before examination. Diffusion MRI data were acquired using all four PEDs, with each PED containing acquisitions of three low *b* values (one *b* = 0 *s*/*mm*^2^, two *b* = 50 *s*/*mm*^2^, two *b* = 300 *s*/*mm*^2^ and 32 *b* = 1100*s*/*mm*^2^images). Acquisitions were performed using a 32-channel head coil, a SENSE factor of 2 with no simultaneous multi-slice, no partial Fourier encoding, and an isotropic resolution of 2*mm* (TE/TR: 92/12875 *ms*). A square field of view was employed for all acquisitions to ensure identical echo train lengths for both AP/PA and RL/LR phase encoding. In addition to dMRI, fat-suppressed T2W turbo spin echo was acquired as a structural image both to complement susceptibility distortion correction and to rigidly align all processed PED dMRI data to a common subject-template space.

The standard vendor-provided phase correction for removing ghost artifacts was applied during image reconstruction.

#### 2.4.1 dMRI processing

The preprocessing of the dMRI data was performed using the TORTOISE software pipeline as detailed in (11). First, the DWIs were corrected for Gibbs ringing artifacts (42) and subsequently for motion and eddy-current distortions (43). For EPI distortions, including susceptibility-induced and concomitant field distortions, a pair of reversed PEDs (11) (i.e., AP^raw^ and PA^raw^, or LR^raw^ and RL^raw^) were used together to generate a processed dataset, denoted as AP, PA, LR and RL. For all PED datasets, the Jacobian modulated signals were output in these corrected images. To provide a common space for longitudinal analysis, the T2W structural image of the first scan session for each subject was manually reoriented to anterior-commissure to posterior -commissure (ACPC) orientation, denoted as T2W^ACPC^. The DWIs for all scan sessions across all PEDs for a subject were rigidly aligned to this ACPC-reoriented structural image.

**Figure 3** demonstrates the integration of synthetic ghost images within dMRI processing pipeline. To capture artifact characteristics at each stage the pipeline, synthetic ghosts were generated in each subject’s native space, as shown in **Fig 2**, and transformed into the final template space, where all PED images were aligned. To transition from native space to template space, all transformations from the processing steps of dMRI data including those for eddy-current correction, EPI distortion correction, and dMRI-to-T2W alignment, were combined into a single warp from native to template space. **Figure 4** displays the overlayed synthetic ghost images on top of the *b* = 0 images before and after dMRI processing. We observed a strong alignment of those synthetic ghost images with abnormal patterns that exist in **Fig. 1**, supporting our hypothesis. Following this, an automatic Otsu thresholding and morphological process (closing) were deployed onto the transformed synthetic images to generate residual ghost maps (*i.e.,* Mask(AP^syn^) and Mask(PA^syn^)).

**Fig. 3.**
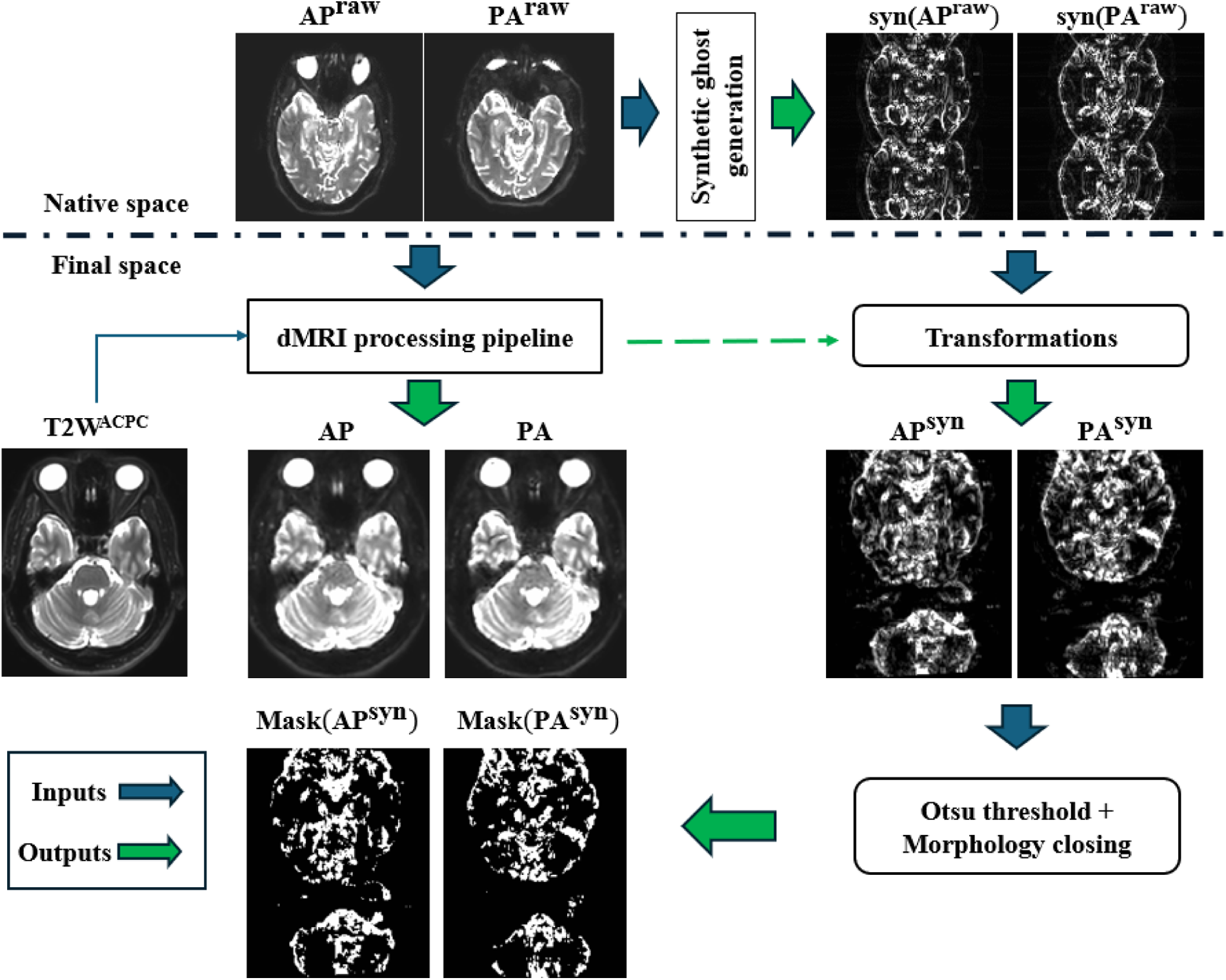
Integration of synthetic ghost images into the dMRI processing pipeline. Synthetic ghost images syn(AP^raw^) and syn(PA^raw^) were generated from unprocessed images AP^raw^ and PA^raw^ in the native space, using the corresponding PED and undersampling factor, R. A pair of reversed PEDs images were input (blue arrow) through the processing pipeline with EPI distortion correction, and rigidly aligned to the T2W^ACPC^, resulting in AP and PA. The transformations in dMRI processing were warped and applied to these synthetic ghosts (green arrow), resulting in AP^syn^ and PA^syn^. After alignment with the processed data, the binary ghost maps were generated for subsequent variance-based analysis. A similar process was also applied to LR^raw^ and RL^raw^ images.

**Fig. 4.**
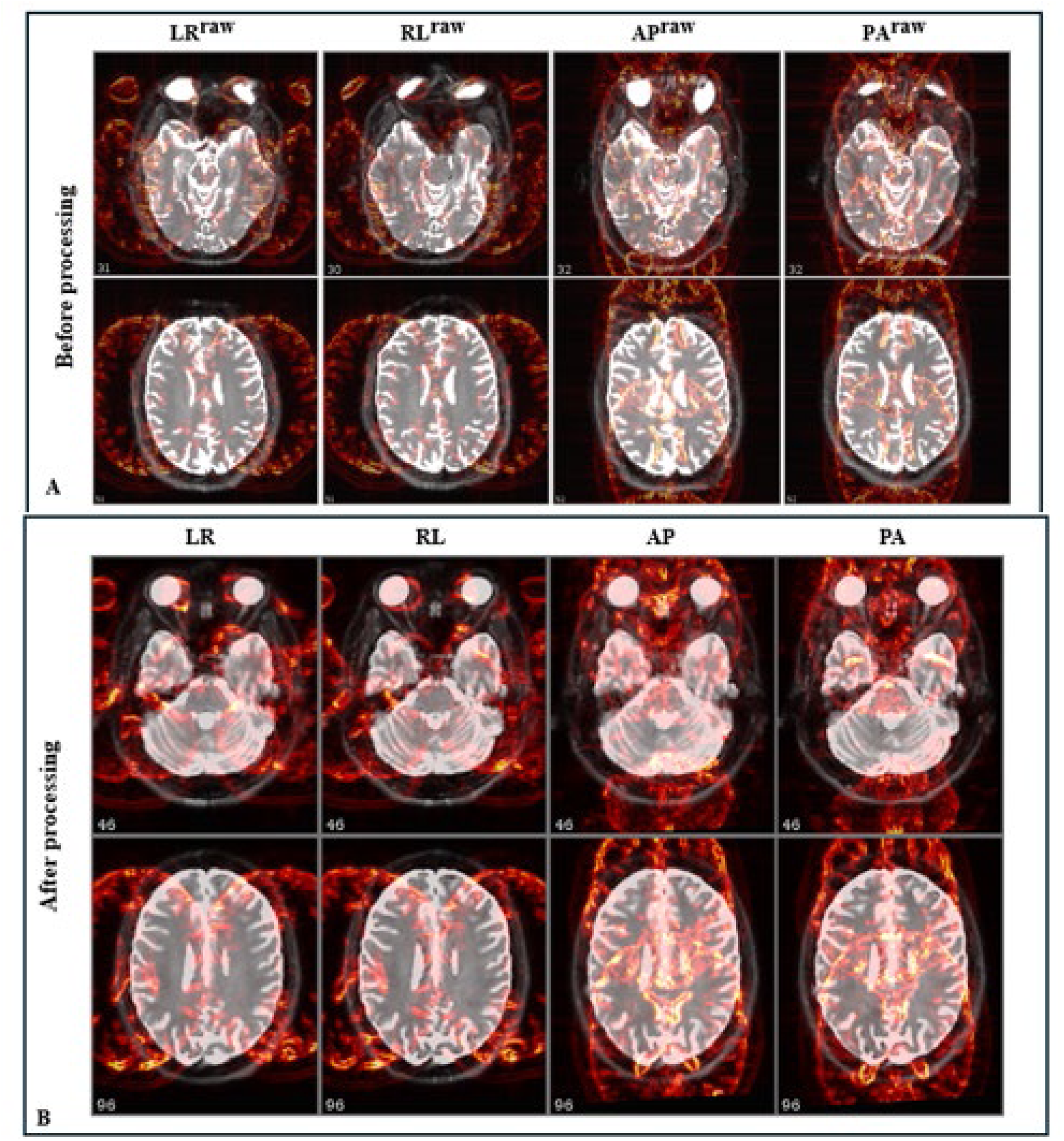
Overlaying synthetic ghost images (red) on top of ***b* = 0 *s*/*mm*^2^** images before [LR^raw^, RL^raw^, AP^raw^, PA^raw^] and after processing [LR, RL, AP, PA] of the in vivo data. Panel A displays the synthetic ghost images generated for each PED in their native spaces. Residual artifacts in LR^raw^ and RL^raw^ images before processing step are hardly visible, while the residual artifacts impact on the AP^raw^, and PA^raw^ images are aligned with the synthetic residual artifacts’ images generated by proposed method. Panel B shows the transformed synthetic ghost images into the final space. Since all images across PEDs has been aligned to the same space, the example slices are consistent. The synthetic residual ghost images in red aligned with the abnormal patterns that exist in those processed images. Slices across PEDs are manually selected to obtain similar structures.

#### 2.4.2 Reproducibility evaluation

Residual artifacts were identified by comparing differences across all PEDs, similar to spotting subtle differences between images, as indicated by yellow arrows in post-processed images shown **Fig. 1**. For quantitative validation, we examined the longitudinal variability for each subject. Since these scans were acquired in a short timeframe, we assume minimal biological changes. Therefore, any disparities observed between longitudinal scans can be attributed to experimental variability and processing differences.

We evaluated the correction performance on the in-vivo dataset by analyzing the longitudinal variability of each subject’s data across multiple scans. For each scan, we computed Fractional Anisotropy (FA) and Mean Diffusivity (MD) maps from the processed datasets for each PED, including LR, RL, AP, PA and the combined dataset *APLR*_*C*0_, *APLR*_*C*1_, and *APLR*_*C*2_. Here, *APLR*_*C*0_ represents the dataset that is obtained by concatenating all the DWIs from [LR, RL, AP, PA] as performed in previous work (12). Additionally, *APLR*_*C*1_, and *APLR*_*C*2_ are the modified version of *APLR*_*C*0_ where the artifactual *b* = 0 *s*/*mm*^2^images were replaced by the corrected *C*_1_(*b* = 0) and *C*_2_(*b* = 0), respectively.

As described in Sec. 2.4.1, each subject scan was rigidly aligned to the T2W^ACPC^ of the first scan to ensure correspondence across scans. Longitudinal variability was assessed by generating voxelwise standard deviation (SD) maps of FA and MD for each subject. We hypothesize that reduced variabilities (i.e., SD values are lower) after the application of our method indicate better artifact removal performance. To assess longitudinal variability at a population level, a DTI template using diffusion tensor data from all subjects was first created (44). The transformations mapping each subject to the template space were stored, and applied to the individual subject SD, aligning them with population template. Once aligned, an average map is computed across all subjects. Examining this average map allows us to observe topological patterns, identifying specific regions that are stable and reliable across cohorts and highlighting the impact of artifact corrections.

## 3. Results

### 3.1 Detection and correction in simulated data

**Figure 5** displays the artifact detection results corresponding to the R=1 simulated sample set illustrated in **Fig. S3**. In this simulation sample, the detection rate was high, as illustrated by the predominance of yellow labels on these images, especially in Case 1 and Case 2. This demonstrated that the proposed method could detect artifacts that cause inhomogeneity across PEDs at the voxel level, without prior information of which images were affected. However, as the number of artifact-affected images increased in Case 3 and Case 4, this resulted in an increase in the number of anomalous values in data across the PEDs in some voxels, which weakened the VBA statistical assessment. As a result, the occurrence of FN rose, resulting in a decrease in Dice scores across artifact-affected PED images. Artifact detection results for the R=2 simulated sample set are displayed in **Fig. S4**. With the increased number of ghosts with R=2, the detection rate was lower as more artifacts overlapped with each other across PED images.

**Fig. 5.**
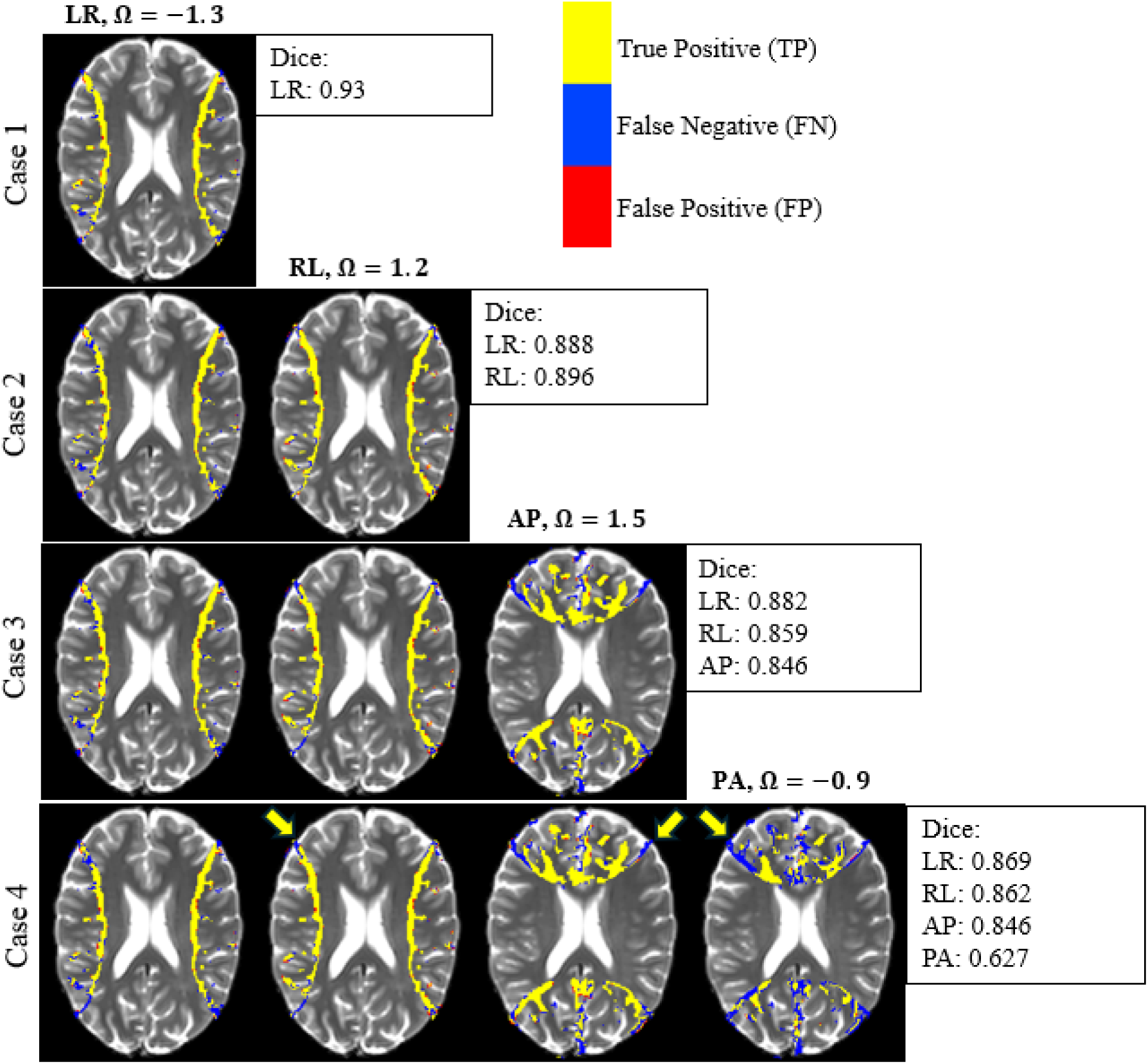
Artifact detection performance on a simulated dataset across all cases. This dataset mimics fully sampled k-space EPI data (R=1) where residual artifacts appear at N/2 positions along the PED of the image. The intensity of artifacts of each PED, controlled by ***Ω***, were ***Ω*_*LR*_ = −1.3**, ***Ω*_*RL*_ = 1.2**, ***Ω*_*AP*_ = 1.5**, and ***Ω*_*PA*_ = −0.9**, remaining constant across all cases. The yellow, red, and blue pixels illustrate the true-positives (TP), false-positives (FP), and false-negatives (FN) of the detection method in artifact-affected images. Dice scores for each case are displayed to the right of the images, quantifying the detection accuracy. The yellow arrows highlight the voxels where artifacts from more than two PEDs (*e.g.* artifacts of LR and AP) overlapped with each other, resulting in increased FN pixels.

To quantify the results of detection under different scenarios, **Table 1** presents the overall mean and standard deviation of Dice, Precision, and Recall scores of artifact detections over 100 realizations for each scenario. A gradual decline, from Case 1 to Case 4, in Dice score from both R=1 and R=2 is associated with the number of artifact-affected images across all PEDs. The Precision scores are high across all cases and conditions, indicating the true-positive rate was close to 0.9, reflecting the detection method was reliable in identifying the actual artifact voxels. The Dice and Recall scores decreased due to the penalty for false negatives, which were exacerbated by the increased overlap of ghost artifacts in the images. Our results suggest that the detection method maintains high precision but suffers in recall and Dice score as the complexity (number of artifact-affected images and R-factor) increases.

**Table 1:** Mean (SD) of performance metrics including Dice, Precision and Recall scores for the detected residual artifacts in the simulated dataset. The metrics were computed over 100 realizations of Ω ghost intensities. The Precision scores indicate that the detection method was highly reliable in all cases, with 90% of the positive predictions being correct. However, as the number of artifact-affected images increases and more artifacts overlap across PED images, the number of FN rise, and the Dice and Recall scores drop significantly.

| Table 1 | R=1 |  |  | R=2 |  |  |
| --- | --- | --- | --- | --- | --- | --- |
|  | Dice | Precision | Recall | Dice | Precision | Recall |
| Case 1 | 0.852<br>(0.049) | 0.933<br>(0.020) | 0.787<br>(0.082) | 0.807<br>(0.023) | 0.897<br>(0.030) | 0.736<br>(0.052) |
| Case 2 | 0.828<br>(0.078) | 0.934<br>(0.013) | 0.743<br>(0.070) | 0.771<br>(0.063) | 0.903<br>(0.021) | 0.678<br>(0.049) |
| Case 3 | 0.799<br>(0.098) | 0.933<br>(0.011) | 0.701<br>(0.052) | 0.723<br>(0.093) | 0.905<br>(0.017) | 0.613<br>(0.040) |
| Case 4 | 0.789<br>(0.097) | 0.934<br>(0.009) | 0.681<br>(0.039) | 0.690<br>(0.099) | 0.904<br>(0.012) | 0.568<br>(0.033) |

**Figure 6** provides a visual and quantitative evaluation of the correction techniques for the R=2, Case 2 simulated data, assuming that residual artifacts only exist in the LR and AP PEDs. Three different correction methods, averaging, CorrV1 and CorrV2, were applied. The results are shown in the second row respectively as *C*_0_, *C*_1_, and *C*_2_; and the %*Err* maps computed between those corrected images against the ideal image are shown in the third row. Although residual ghost artifacts in the *C*_0_ image were suppressed compared to the LR and AP images, they were not completely removed as indicated by the yellow arrows. The %*Err*_*C*1_ and %*Err*_*C*2_ exhibit lower errors than %*Err*_*C*0_ image, particularly in areas where the ghost artifacts of two different PEDs did not overlap. In the voxels where artifacts in LR and AP overlapped, the residual artifacts were not completely suppressed and were observed in a higher %*Err*_*C*1_. However, the average of %*Err* rates are 0.54% and 0.28% for corrected images *C*_1_, and *C*_2_ respectively, which are markedly less than the average %*Err* error observed in the *C*_0_ image of 2.42%. This underscores the effectiveness of the CorrV1 and CorrV2 methods in minimizing artifacts, especially CorrV2.

**Fig. 6.**
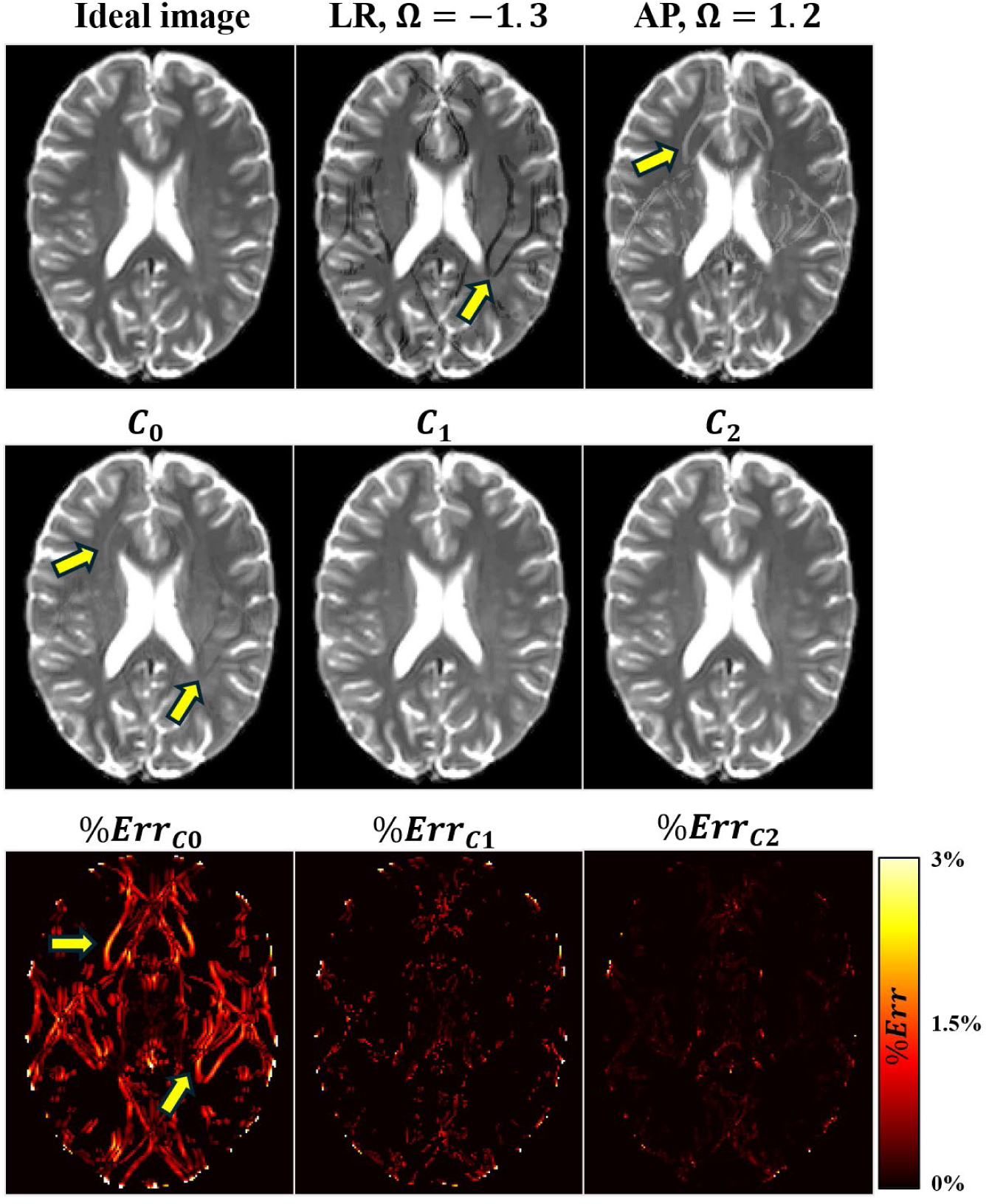
Comparison of correction methods against the ideal image for the parallel imaging factor R=2, Case 2 (two artifactual images) simulated data. In this simulation sample set, only LR and AP images had residual artifacts, with *Ω*-values of −1.3 and 1.2 respectively. The first row displays from left to right, the ideal image, the artifact-affected images, LR and AP. Other PED images such as RL and PA images are not shown, as they are identical to the ideal image. The second row displays the mean image (***C*_0_**) derived from averaging all four PED images, and the images corrected using CorrV1 (***C*_1_**) and CorrV2 (***C*_2_**) approaches respectively. Yellow arrows indicate visible residual artifacts in the LR and AP images in the ***C*_0_** image, which are not present in the C1 and C2 images. The third row showcases the %***Err*** images, illustrating the percentage error when comparing the ***C*_0_** and corrected images (***C*_1_ and *C*_2_**) against the ideal image. The images obtained from both correction methods outperformed the averaging approach. ***C*_1_** and ***C*_2_** images show no visible residual artifacts compared to the ***C*_0_**, and the %***Err*** of these corrected images was also lower than the %***Err*** of the ***C*_0_**.

**Figure 7** presents a comparative analysis of the average (*C*_0_), CorrV1 (*C*_1_) and CorrV2 (*C*_2_) correction methods across two configurations R=1, and R=2, using the %*Err* and SSIM metrics over 100 realizations of simulated data. The results highlight a clear superiority of the correction methods over the averaging approach, with both *C*_1_ and *C*_2_ significantly reducing errors (%*Err*) and improving structural similarity as indicated by increased SSIM. Notably, *C*_2_ was better than both *C*_0_ and *C*_1_ images with fewer overall errors. However, in case 4, the correction performance of CorrV1 was not as effective as the averaging method. This was due to the incomplete removal of error datapoints in regions with overlapping artifacts across all PEDs, resulting in inaccurate estimation. As a result, %*Err* of *C*_1_ in case 4 was higher than *C*_0_.

**Fig. 7.**
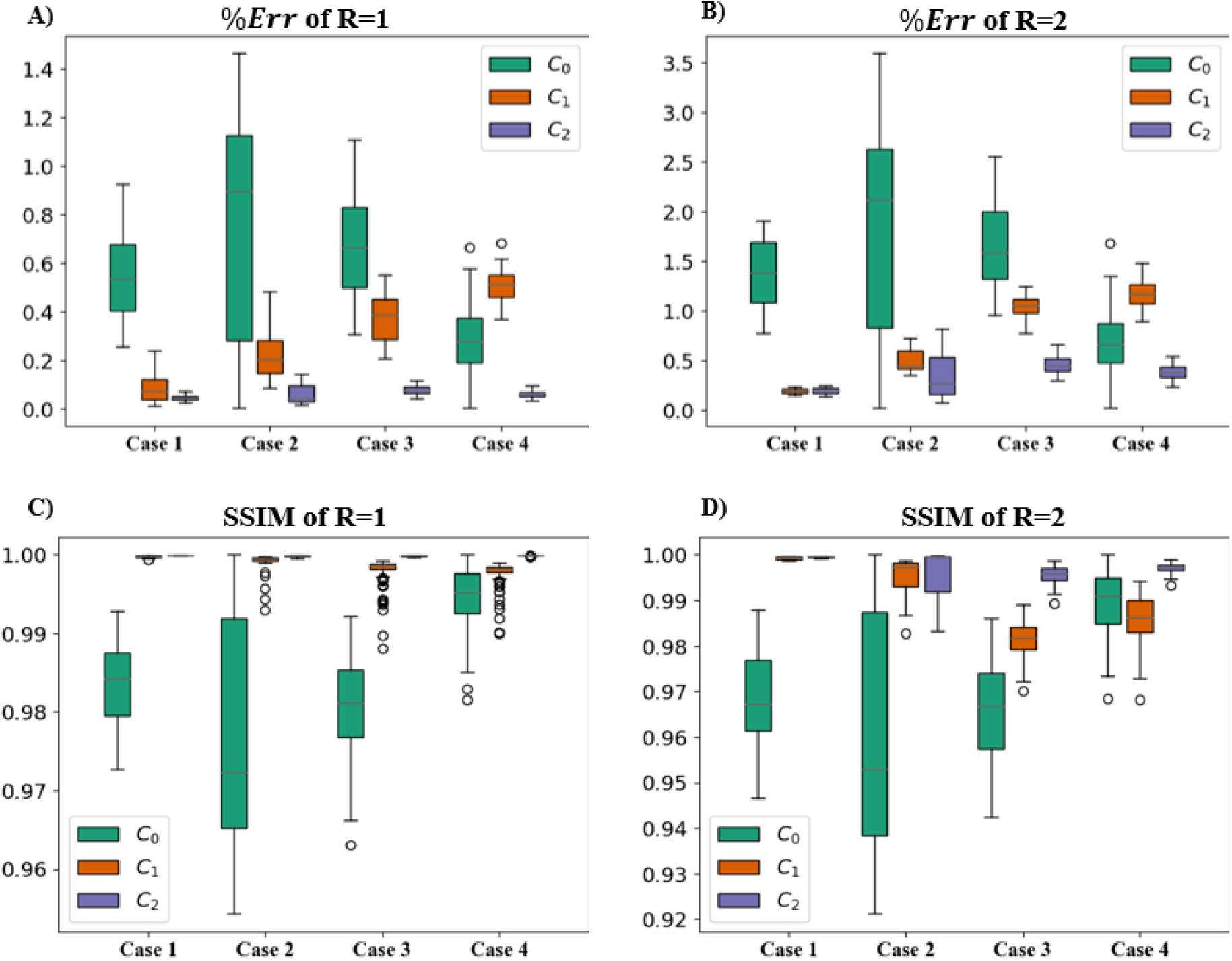
Performance comparison of correction methods across R=1 and R=2 simulated data in all cases. The figure presents an image quality comparison of ***C*_0_**, ***C*_1_**, and ***C*_2_** based on 100 realizations of simulated data under two configurations: R=1 (A, C), and R=2 (B, D) across four cases. Box plots for %***Err*** (top row) and SSIM (bottom row) illustrate the image quality improvements from CorrV1 (***C*_1_**) and CorrV2 (***C*_2_**) compared to the average images (***C*_0_**). The results demonstrated a reduction in errors and an increase in structural similarity, indicating enhanced image quality.

### 3.2 Detection and correction of in-vivo data

**Figure 8** displays results of automatic artifact detection using our proposed method on the corresponding slices shown in **Fig. 1**. The VBA measurements (i.e. %***X***_*jj*=*PED*_) exceeding the heuristic threshold are overlayed on the slices. At locations where the residual artifacts are prominently different across PED images the VBA measure is considerably large (≥ 50%). Our method successfully identifies these artifacts, aligning with the visually observed residuals, therefore validating the model’s effectiveness in automated artifacts detection. Additional examples of artifact detection results from different slices of the same representative subject are shown in **Fig. S5.**

**Fig. 8.**
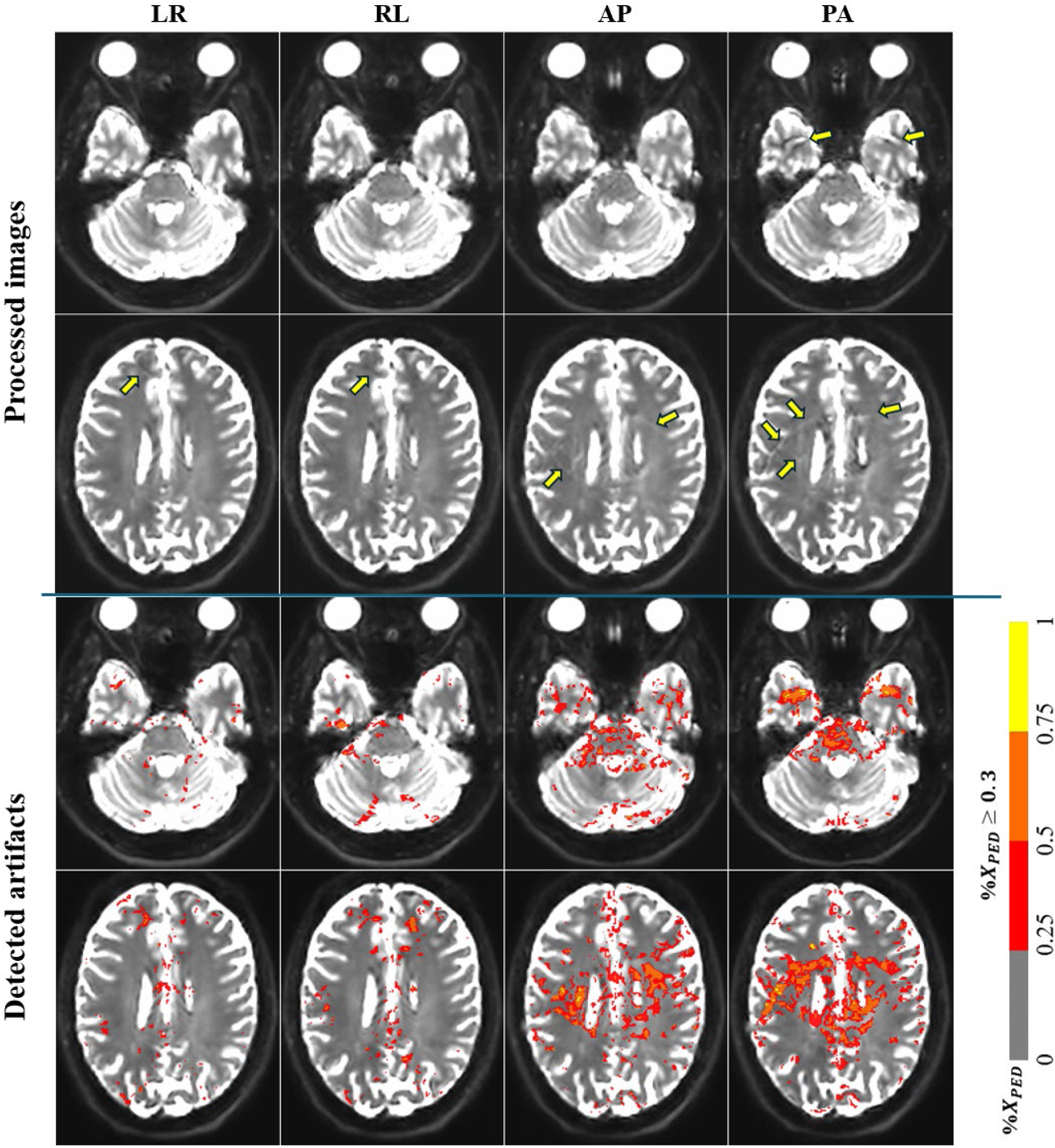
Detected residual ghost artifacts in the processed ***b* = 0** images across LR, RL, AP, and PA from selected slices in Fig. 1. The images are shown twice, with and without the artifact detection overlay, for the reader’s convenience. The detected artifacts, based on the percentage of variance from each PED, exceeding the heuristic threshold (30%), %***X*_*PED*_**, are illustrated using an overlaid segment-colormap. The proposed method was able to detect differences across PEDs as residual ghost artifacts.

**Figure 9** shows the correction results for slices displayed in **Fig. 8**. We observed the remains of residual artifacts in the *C*_0_ image, although they are barely visible. On the other hand, while both CorrV1 and CorrV2 methods effectively reduced artifact appearances, C1 images exhibit patchiness due to intensity differences across PEDs, while C2 images were smoother, demonstrating superior refinement in artifact mitigation with this dataset.

**Fig. 9.**
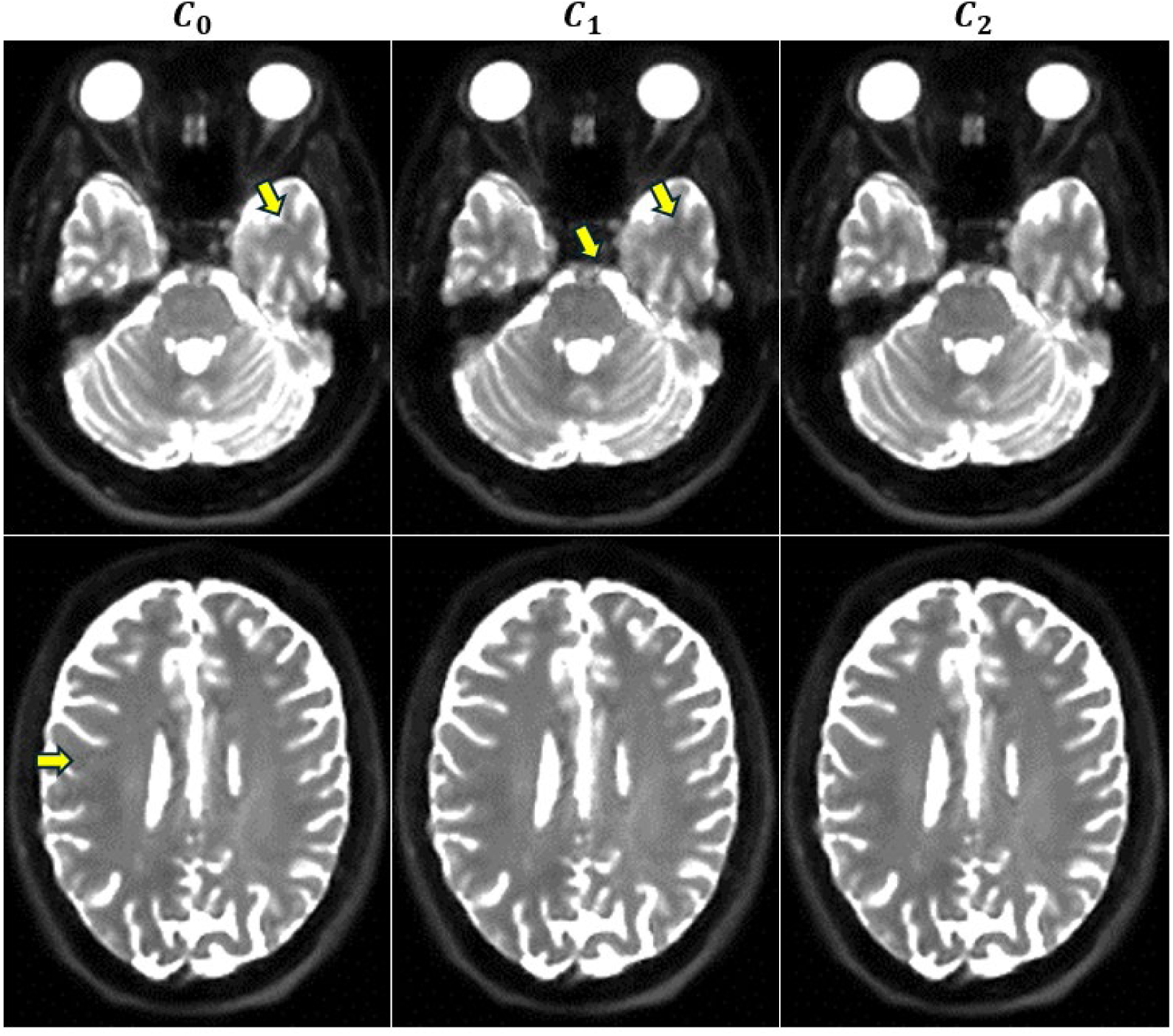
Corrected images from averaging (***C*_0_**), CorrV1 (***C*_1_**), and CorrV2 (***C*_2_**). Although residual artifacts are suppressed and hardly visible in the C_0_ image, the ***C*_1_** and ***C*_2_** images show better suppression in the areas with residual ghost artifacts as indicated by the yellow arrows in C_0_. The ***C*_1_** images exhibit some pixelated values (patchiness), indicated by the yellow arrows, due to the intensity differences across PEDs, while the ***C*_2_** images appear smoother and more uniform.

To quantify the correction results at a population level, **Fig. 10** displays the averaged SD maps of FA and MD in the population template space, specifically for the *AP*, *APLR*_*C*0_, *APLR*_*C*1_, and *APLR*_*C*2_ datasets. In this figure, bright regions correspond to areas of lower reproducibility, with higher variation across scans, while darker regions indicate areas of higher stability. In the 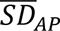 map, residual eye ghosts cause instability, as shown by the bright regions in yellow circle. Similarly, in the same area, in 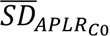 (indicated by yellow arrow), residual artifacts from *AP* are carried into *APLR*_*C*0_ since it represents an average of all four PEDs.

**Fig. 10.**
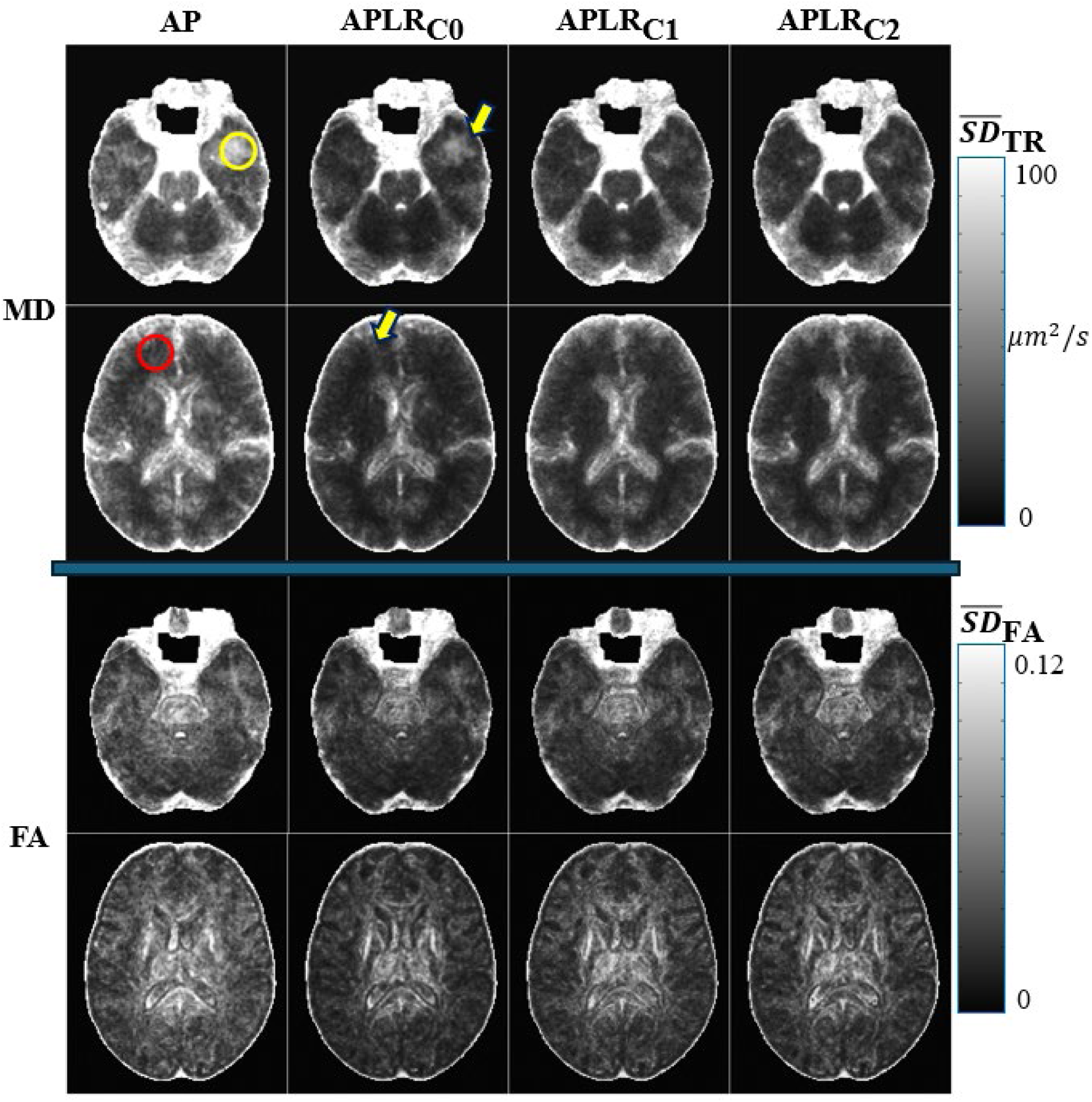
Population average of standard deviation maps at two slice levels for FA and MD. Columns represent images generated from different processing methods (*i.e.* ***AP*, *APLR*_*C*0_, *APLR*_*C*1_**, and ***APLR*_*C*2_**). The bright regions indicate higher average variation, while the darker regions denote lower overall variation. The arrows indicate the remnants of residual artifacts. The red and yellow circles indicate the regions of interest which are the temporal lobe (ROI-1), and the midbrain regions (ROI-2) respectively. The increased overall variability in the AP dataset originates from the number of volumes used in the diffusion tensor estimation, which was half of the other datasets. This reduced number of datapoints in the estimation caused an average increase of sqrt(2) in the computed standard deviations maps.

Compared with *APLR*_*C*0_,the *APLR*_*C*1_ and *APLR*_*C*2_ datasets, which include corrected *b* = 0 images with artifact removal, demonstrate marked improvements in stability. Another key point is that correction effects were less evident on average SD map of FA than MD. This difference arises because MD measure the total diffusivity, making it more influenced by absolute signal intensities, (such as *b* = 0 images), while FA reflects the directional dependence of diffusivity which is more sensitive to relative differences between diffusivities in different DWIs. **Figure S6** and **Figure S7** illustrate these differences on the average SD maps of MD and FA images from a representative subject.

Visual comparisons across PEDs from the in vivo findings highlight the detection of residual artifacts across PEDs. Increasing the heuristic threshold in VBA should limit sensitivity to variance of artifacts across PEDs. Without ground-truth images, quantitative comparison using SD maps revealed that the regional effects of residual ghost artifacts on longitudinal reproducibility were further reduced by our correction methods, specifically in MD images. For example, **Fig. 10** and **Fig. 11**, in the temporal lobes, which are affected by ghosting from the eyes in AP-encoded data, the *APLR*_*C*0_ demonstrated a 50% reduction in variability (in agreement with the Irfanoglu et al. (12)). On the other hand, *APLR*_*C*1_ and *APLR*_*C*2_ both achieved an additional 30% decrease in variability compared to *APLR*_*C*0_.

**Fig. 11.**
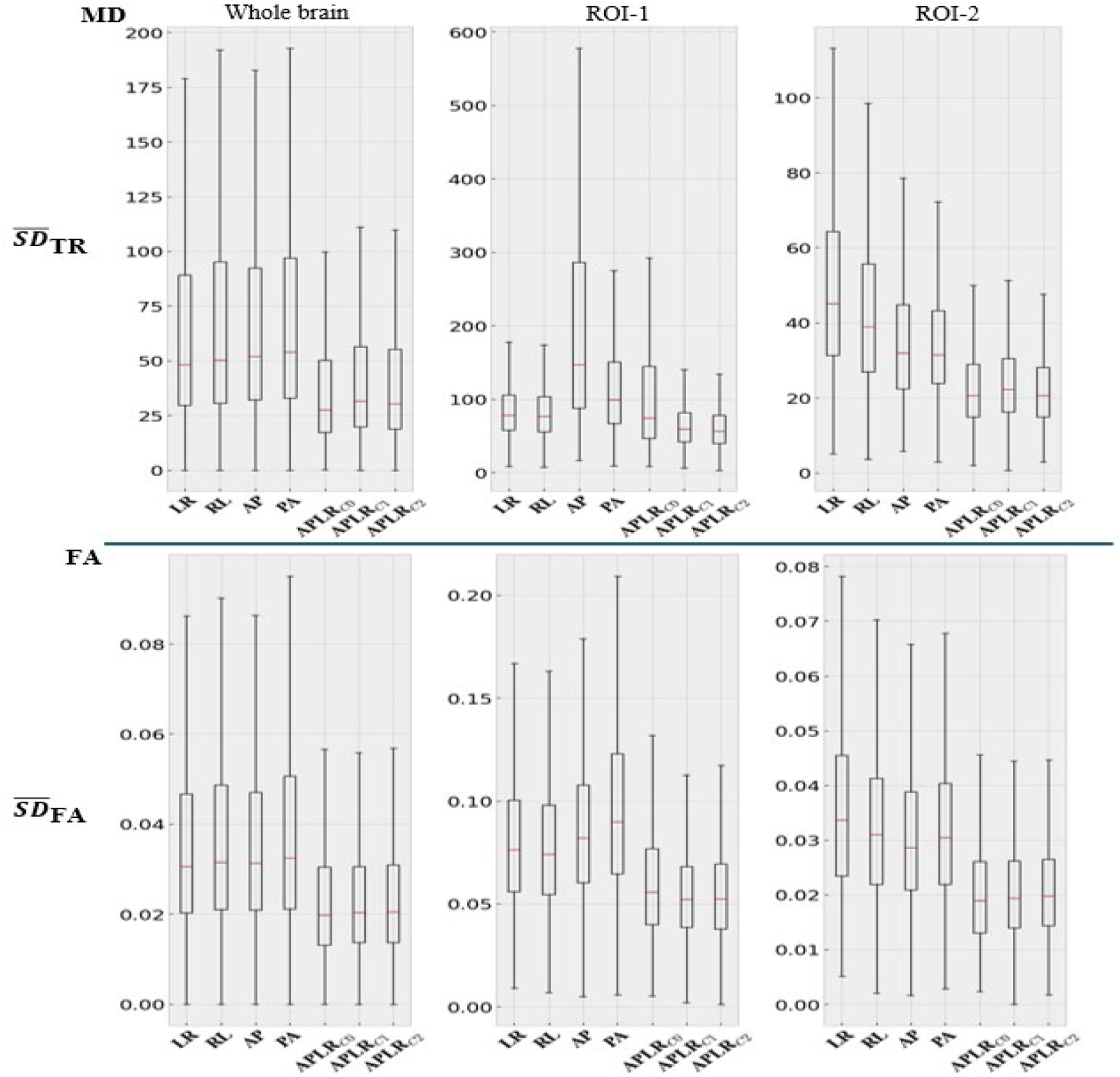
The ROI-wise distribution of longitudinal average SD values. The error bars represent the standard deviation of the average values across the population. Moving from single phase-encoding to four-way phase encoding significantly improves reproducibility for both metrics. The proposed correction strategies CorrV1 and CorrV2 clearly improve the longitudinal reproducibility in artifact-affected regions (ROI-1 and ROI-2) and do not negatively affect the reproducibility within the whole brain in a significant manner.

**Figure 11** provides a quantitative comparison of reproducibility across the entire population for each processed dataset. Focusing on three specific regions of interest (ROIs): whole brain, the temporal lobe (ROI-1, indicated by a yellow circle in the first row of **Fig. 10**), and midbrain (ROI-2, indicated by a red circle in the second row of **Fig. 10)**. In the whole-brain analysis, a slightly increased variability is observed in the datasets processed through CorrV1 and CorrV2 methods, compared to *APLR*_*C*0_ for both MD and FA. This can be attributed to the lower number of *b* = 0 images used for the estimation of the diffusion tensor, which was reduced by the replacement of corrected *b* = 0 images in *APLR*_*C*1_ and *APLR*_*C*2_ sets. However, improvements were still significant compared to the cases with no correction, *APLR*_*C*0_. For the two regions of interest, which are predominantly affected by artifacts, the proposed correction methods enhanced reproducibility. For ROI-1, which was affected by ghosting of the eyes in AP and PA PEDs, the variability levels were significantly reduced bringing them in line with the stability observed in the LR and RL PEDs. A similar improvement was also observed for ROI-2.

## 4. Discussion

Acquiring four-PEDs data is not a conventional approach for dMRI or other imaging modalities, as it requires manual adjustments to ensure identical acquisition parameters across PEDs, such as FOV, echo time, repetition time, and diffusion times. However, a previous study (12) has demonstrated the benefits of using four-PED data in improvement of reproducibility, and here we presented an expanded application to enhance data reliability through residual artifacts removal. The core concept of this approach is to reduce significant artifact components that could affect quantitative analysis. While our method focused on residual ghost artifacts, as an illustrative artifact type, we believe it could potentially be adapted to other types of artifacts, such as chemical shift and parallel imaging artifacts, by adjusting a synthetic model to align with specific artifact characteristics. For instance, EPI ghost artifacts can be modeled based on odd/even echo separation, whereas chemical shift artifacts are influenced by water-fat differences and phase-encoding bandwidth.

Although dMRI served as the primary imaging modality in this study, this method may also be applicable to other EPI-related modalities, such as fMRI, which consists of non-diffusion-weighted volumes over time. This setup could benefit VBA analysis by increasing the sample size and enhancing artifact detection robustness.

To our knowledge, this is the first study to examine the impact of artifacts throughout the imaging pipeline using four-PED data. However, there are some limitations. First, our focus was restricted to non-diffusion-weighted (*b* = 0 *s*/*mm*^2^) images, due to the significant impact of eddy-current effects on diffusion-weighted images. Eddy current, induced by rapid switching of diffusion gradients, can distort images by introducing spatial misalignment and intensity variations. These distortions can cause geometric warping, signal dropout and shifts in structure locations within the image. Consequently, eddy-current effects can complicate the VBA method, as they affect not only the true-anatomical features but also the artifacts, making it difficult to separate real signal variations from artifact-induced changes. Moreover, as diffusion weighting increases, ghost artifacts become less prominent, reducing the contrast between PEDs and resulting in insufficient information for effective comparison. Second, while our study focused on data acquired with SENSE (R=2), single-shot and full k-space coverage, variations in these parameters such as switching to multi-shot or using partial Fourier can alter artifact manifestations, necessitating different models to accurately map residual artifacts.

Finally, the underlying assumption of our approach, which states that a voxel is artifactual in at most two PEDs, may not hold when acquiring images with higher parallel imaging factors (R ≥ 2), which causes increased aliasing. Therefore, a voxel in a given PED may be affected by one ghost, while the same voxel in a different PED image may be affected by different ghosts (or other artifacts). This behavior was observed in a small number of voxels in simulated data (case 4) and in-vivo result **Fig. S5B**, where the yellow arrow highlights the misidentification of artifact detection)

Our future work will pursue two main goals. First, we plan to investigate residual artifacts directly from actual k-space data during the acquisition/reconstruction stage, if applicable, rather than relying on synthetic models to test and improve future MRI systems. Second, we aim to create a database consisting of a large population of four-way encoded data along with corresponding detected and corrected results to train deep neural networks. These networks could then be applied to other single PED data as an additional retrospective approach for enhancing the data reliability.

## 5. Conclusion

This study leveraged four-way PED data to address the challenge of detecting and correcting residual EPI ghost artifacts in dMRI data. By developing a retrospective, image-based correction strategy, we filled a gap in existing dMRI processing pipelines, providing a method that integrates well with current practices. The potential application of this method to other EPI modalities such as fMRI, or its use with less than 4-way phase-encoding acquisitions are promising directions that can further improve the impact of this work.

## Code and Data Availability

The data used in this experiment were analyzed retrospectively. The data are not publicly available but can be provided on special request. Please contact the last author if you would like access to the data. The code for the simulation data is available at: https://github.com/15thai/Residual-Artifact-Detection-with-Four-PEDs-data

## Acknowledgments

The authors would like to thank Dr. Neville Gai for advice on Nyquist ghost in in-vivo data and simulated data, as well as the NIH Fellows Editorial Board and Lindsay Walker for reviewing and editing the manuscript.

This work has been funded by the National Institute of Biomedical and Bioengineering Intramural program. The contents of this work do not necessarily reflect the position or the policy of the government, and no official endorsement should be inferred.

**Anh Thai** is currently a Ph.D. student at Catholic University of America, where she also received her B.S. and M.S. in Electrical Engineering in 2015 and 2017. Her current research interests include machine learning and advanced image processing methods for medical applications.

**Lin-Ching Chang Ph.D.** received her Ph.D. in Computer Science from George Washington University and serves as a full professor of Computer Science at the Catholic University of America, where she also direct the Data Analytic program. Her research focuses on machine learning, data science, medical image processing, and biomedical data analytics. Dr. Change is known for her interdisciplinary approach, advancing computational methods to address complex challenges in healthcare, data analysis and AI.

**Carlo Pierpaoli M.D., Ph.D.** is a senior investigator of the Quantitative Medical Imaging (QMI) Lab at the NIBIB, NIH. He received his M.D. in neurology and a Ph.D. in neuroscience at the University of Milan, Italy. He has spent most of his scientific career at the NIH, where he is also well-known for his contributions in the field of diffusion MRI applied to brain studies. He is a fellow of the International Society of Magnetic Resonance in Medicine and received the NIH award of Merit for performing the first diffusion tensor imaging studies of human brain.

**M. Okan Irfanoglu Ph.D**. is a staff scientist at the QMI Lab at NIBIB, NIH. He received his PhD in Computer Science and Engineering department of Ohio State University. He has been designing and developing the TORTOISE software, which aim to extract accurate and reproducible biomarkers from data acquired with MRI. He is an expert in image and signal processing, who has contributed to the field of diffusion MRI for over 16 years.

