## Supporting Figures for "Exploiting Four-way Phase-encoding Benefits for Robust Detection and Correction of EPI Artifacts: Application to Residual Ghosts in Diffusion MRI"

### Supporting Information

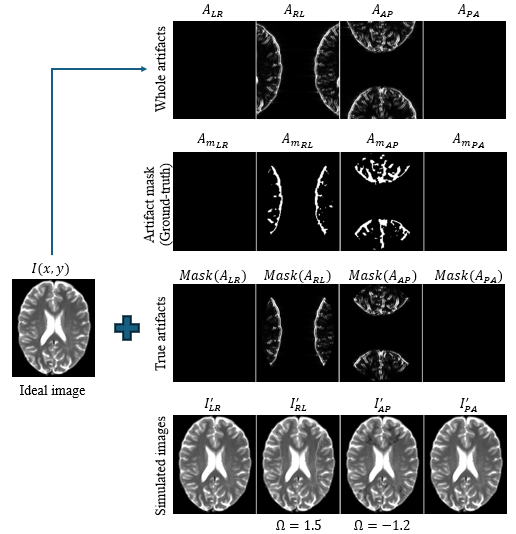

**Fig. S1** The schematic demonstration for the simulated residual artifact images of R=1, Case 2 scenario. A representative slice of T2W image is used as the ideal image. A residual artifact image is generated based on Equation (9), with randomized $\Omega$; in this case, $\Omega_{RL}=1.2$, $\Omega_{AP}= -1.3$, while $\Omega_{LR}= \Omega_{PA}=0.0$(i.e. no artifact). Focusing on residual artifacts within the brain region, artifact mask images are produced to label the artifacts in the region of interest, and to serve as the ground truth mask for the detection validation. At the bottom row, the final simulated images with residual artifacts are shown as described in Equation (11).

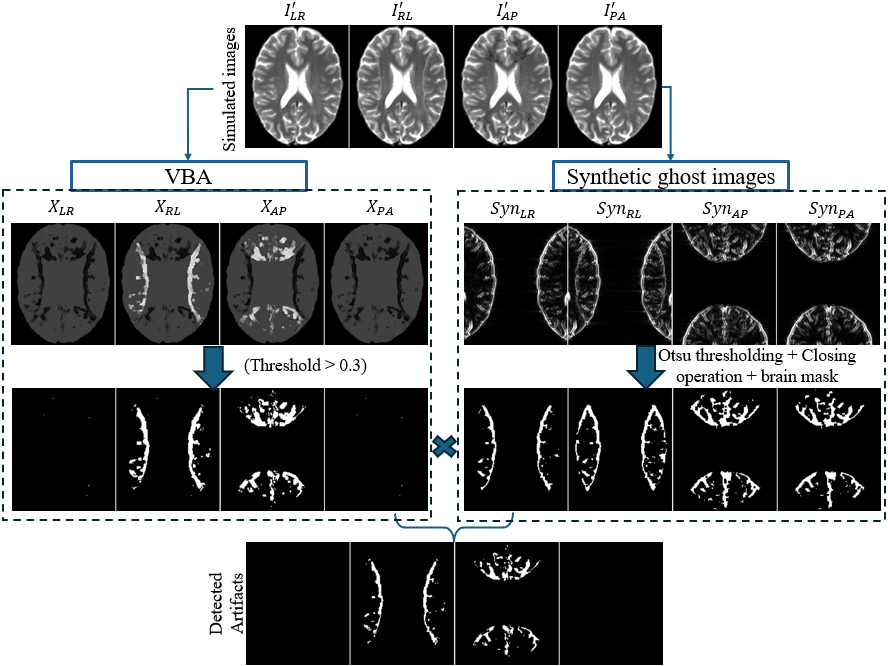

Fig. S2 Demonstration of artifact detection on simulated data, R=1, Case 2 scenario. Both synthetic ghost image generation and VBA are applied to simulated images across PEDs. Outputs from VBA and synthetic image generation are masked to focus on the brain region. Results from VBA are thresholded at 0.3, while Otsu thresholding and closing operations are applied on the synthetic images to label potential residual artifact regions. By combining binary masks from both VBA and synthetic ghost images across PEDs, the arbitrarily introduced simulated artifacts are detected.

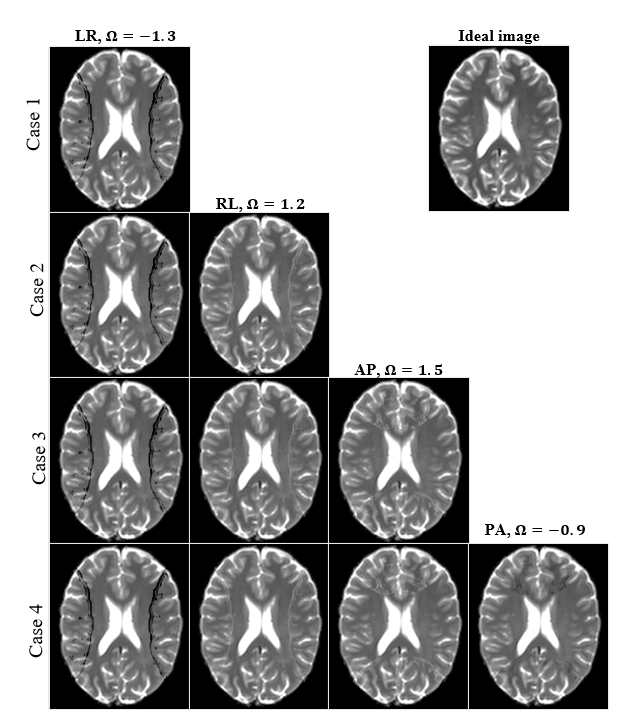

Fig. S3 Example of simulated artifacts of R=1 sample set across different cases. The ideal image was displayed on the right corner of the figure. From the top to bottom rows, the number of artifact-affected images increase one by one for each case, in the order from LR, RL, AP and then PA respectively. To mimic the unknown impact of residual artifacts in practice, the $\Omega$-values were set differently across PEDs, which were $\Omega_{LR}= -1.3$, $\Omega_{RL}=$1.2, $\Omega_{AP}=1.5$, and $\Omega_{PA}= -0.9$. Positive and negative values of $\Omega$ illustrate the hyper- and hypo- effects of residual artifacts, respectively, on the reconstructed images.

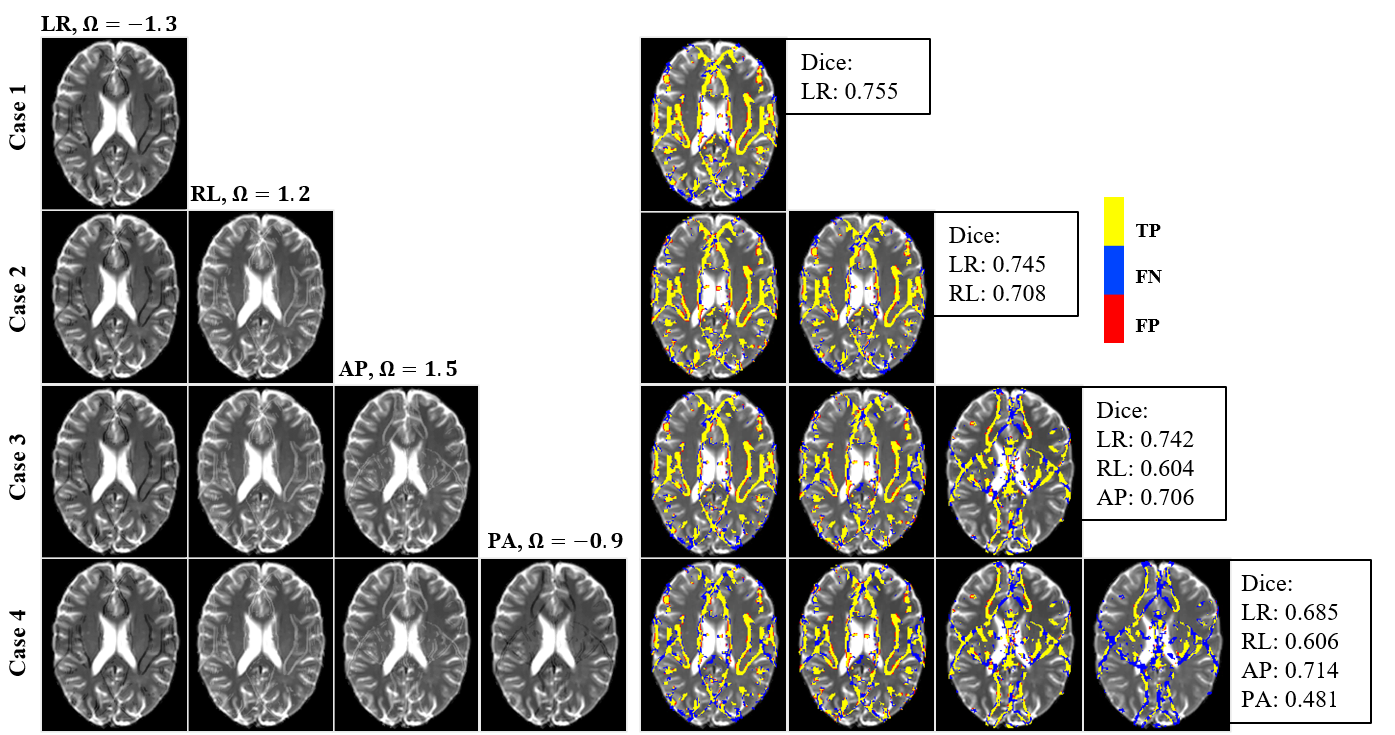

Fig. S4 Display a simulated sample set of R=2 (left) and the corresponding artifact detection results (right) across all cases. In this simulation sample, the residual artifacts appear at N/4 position along the PED of the image. The $\Omega$-values were kept the same as the sample set of R=1, which were $\Omega_{LR}= -1.3$, $\Omega_{RL}=$1.2, $\Omega_{AP}=1.5$, and $\Omega_{PA}= -0.9$. In these cases, the yellow, red, and blue pixels illustrate the true-positive (TP), false-positive (FP), and false-negative (FN) of the detection method in artifact-affected images. The Dice scores for each case in this example were computed.

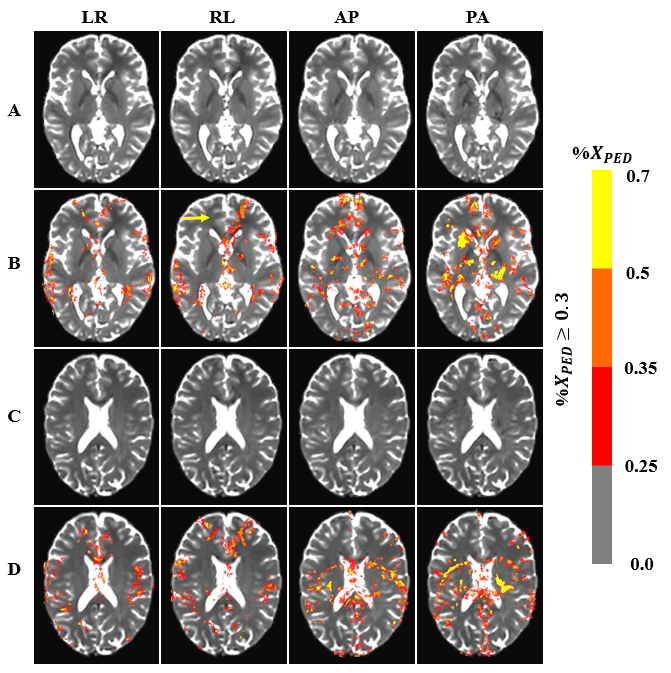

Fig. S5 Another artifact detection results from different slices of a representative subject. The yellow arrow indicates the misidentification of artifacts.

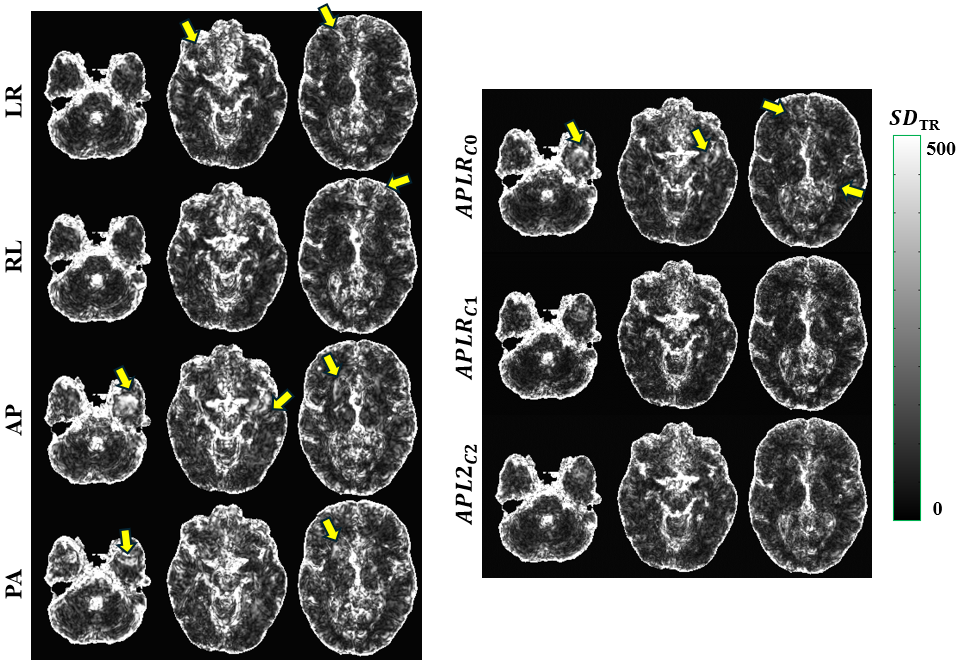

Fig. S6 Reproducibility of MD images of representative subjects across PEDs (LR, RL, AP, and PA) and proposed corrected versions (Mean: $APLR_{C0}$, CorrV1: $APLR_{C1}$, and CorrV2: $APLR_{C2}$). The yellow arrows indicates the impact of residual ghost artifacts across different sets in the longitudinal analysis.

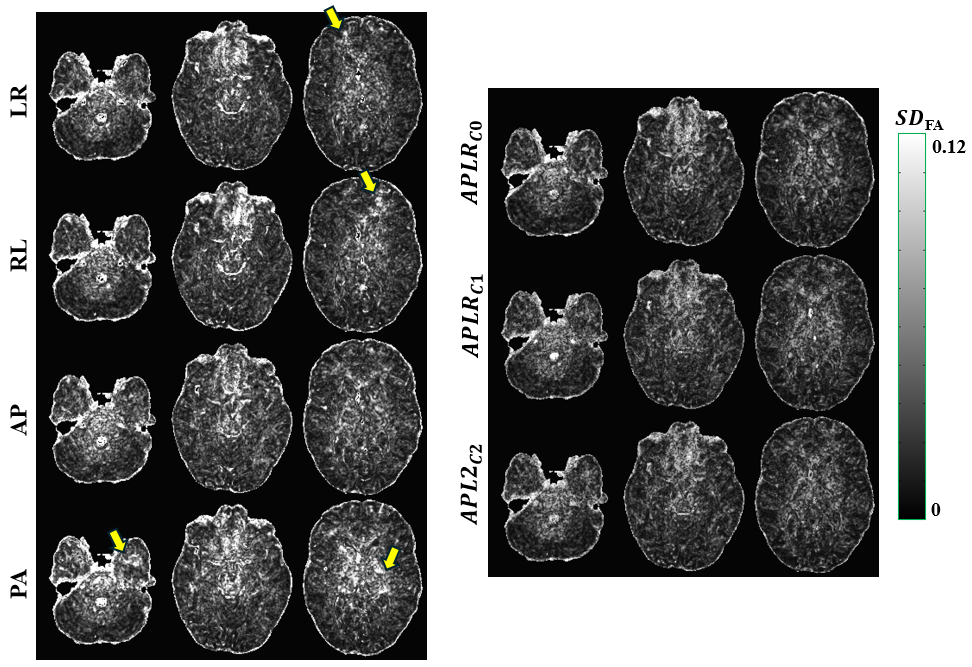

Fig. S7 Reproducibility of FA images of representative subjects across PEDs (LR, RL, AP, and PA) and proposed corrected versions (Mean: $APLR_{C0}$, CorrV1: $APLR_{C1}$, and CorrV2: $APLR_{C2}$). The yellow arrows indicates the impact of residual ghost artifacts across different sets in the longitudinal analysis.
